# Live longitudinal imaging of meningeal cerebrovascular injury and its sequelae in adult zebrafish

**DOI:** 10.1101/2025.11.14.688311

**Authors:** Aurora Kraus, Jean Sebastien Prosper-Santiago, Aleksandra Potapova, John Prevedel, Daniel Castranova, Brant M. Weinstein

**Author notes:** To whom correspondence should be addressed: Brant M. Weinstein Section on Vertebrate Organogenesis Building 6B, Room 4B413, 6 Center Drive, Bethesda, MD 20892.

## Abstract

Nearly 1.4 million people in the United States sustain a traumatic brain injury (TBI) each year, with almost half of those hospitalized for TBI developing long-term disability. For many patients, prolonged bleeding and inflammation from damaged vessels in the meninges result in long-lasting sequelae. Although their injured blood vessels regrow, the site of injury is full of inflammatory immune cells that may influence vascular function. Adult zebrafish have a thin, translucent skull and a mammalian-like meninges that is easily imaged in living animals. We have established a novel adult zebrafish model to investigate vessel-immune cell interactions after meningeal cerebrovascular injury (mCVI). We use carefully calibrated sonication to rupture meningeal blood vessels without breaching the skull or causing damage to the underlying brain. By performing longitudinal live imaging of intubated adult fish we observe vascular regrowth and immune responses to mCVI over time in the same animal with unprecedented resolution allowing measurement of blood flow, dynamics of vessel regrowth, and interactions between individual immune and vascular cells. This newly developed zebrafish model provides a powerful tool for longitudinal live imaging of meningeal immune cell-vascular interactions after cerebrovascular injury, opening the door to new insights into chronic neuroinflammatory disease.

## INTRODUCTION

Although few immune cells infiltrate a healthy brain, it is surrounded by the meninges, a set of immune cell-rich tissue layers that protect vertebrate brains (Coles et al., 2017; Fitzpatrick et al., 2024; Rebejac et al., 2024; Rua and McGavern, 2018; Rustenhoven et al., 2021; Smyth et al., 2024; Xu et al., 2024). Zebrafish, mouse, and human meninges have remarkable anatomical conservation in the construction of the dura mater and leptomeningeal layers, immune cell composition of each layer, and meningeal vascular networks (Absinta et al., 2017; Castranova et al., 2021; Jacob et al., 2022; Louveau et al., 2015; Mecheri et al., 2018; Pietila et al., 2023; Rustenhoven et al., 2021; Shapiro et al., 2021; Venero Galanternik et al., 2025; Walsh et al., 2021). Head trauma often damages meningeal blood vessels, causing meningeal hematomas and meningeal cerebrovascular injuries (mCVI) (Arbona-Lampaya et al., 2025). Head traumas can cause lasting disease pathologies with limited treatment options (Jaja et al., 2018; Maher et al., 2020; Mayer et al., 2002). Acute inflammation facilitates debris clean-up and triggers regeneration, but persistent inflammation correlates with poor long term clinical outcomes (Aertker et al., 2019; Bolte and Lukens, 2021; Fang et al., 2024; Melmed et al., 2021; Ohashi et al., 2023; Russo et al., 2018). Patients with prolonged vascular leakage in the meninges are more likely to have poor clinical outcomes (Hay et al., 2015; Turtzo et al., 2025), and the feedback between meningeal vascular leakage and inflammation is thought to be important in long-term pathology post-mCVI.

A variety of animal models are available to study post-head trauma pathologies, but most of these models focus on neuronal histology, inflammation, and blood brain barrier (BBB) disruption within the brain itself without assessing trauma specific to the surrounding meninges (reviewed in (Bodnar et al., 2019; Chen et al., 2021; Freeman-Jones et al., 2023; Jassam et al., 2017; Siebold et al., 2018; Smith et al., 2021; Zulazmi et al., 2021)). The recent rediscovery of meningeal lymphatic vessels in the dura of vertebrates (Absinta et al., 2017; Castranova et al., 2021; Louveau et al., 2015), has also piqued interest in exploring meningeal functions beyond their traditional role as a barrier for maintaining brain homeostasis (Betsholtz et al., 2024; Salvador et al., 2024). Many types of head injuries can rupture blood vessels in the superficial and highly vascularized meninges, and a few models have been developed with the aim of replicating meningeal vascular pathologies observed in humans. A murine model of subarachnoid hemorrhage involving autologous blood transfer to the magna cisterna to study vascular function after meningeal hematoma revealed that dural lymphatic vessels play a role in debris clearance and modulate neuroinflammation (Chen et al., 2020; Lin et al., 2003; Wang et al., 2023). A murine compression injury model was used for imaging myeloid cell accumulation around meningeal vascular wounds and their role in vascular repair (Roth et al., 2014; Russo et al., 2018). A transcranial ultrasound CVI model was also used in mice to examine how the inflammatory state of the meninges impacts vascular repair (Mastorakos et al., 2021a; Mastorakos et al., 2021b). However, use of each of these models for observation of the meninges involves surgeries and punctures that themselves result in coarse meningeal vascular injuries that cause neuroinflammation.

The structural and functional similarities between the neural, vascular, and immune systems of zebrafish and mammals make the fish well-suited for investigating vascular and immune responses to mCVI (Hotez et al., 2024; Howe et al., 2013; Kenney et al., 2021; Tabor et al., 2019). The availability of innumerable immune cell type-specific transgenic reporter lines has made it possible to carry out live imaging of immune cell dynamics, including new insights into neutrophil migration after injury in larval zebrafish (de Sena-Tomas et al., 2024; Mathias et al., 2006). Longitudinal high-resolution time-lapse imaging of immune cells is possible in adult zebrafish using a non-surgical intubation technique (Castranova et al., 2022; Greenspan et al., 2024). Using this technique, immune cells and vessels can be clearly imaged through the thin, translucent skull of the adult zebrafish, making it possible to use intubation to carry out high-resolution non-invasive live time-lapse and/or longitudinal imaging. As noted above, zebrafish have mammalian-like meninges include both dura mater and leptomeningeal layers, which have been shown to contain cells similar to those found in mammals (Venero Galanternik et al., 2025). As a longstanding model for development and neuroscience, zebrafish also offer a wide array of genetic tools and cell-specific reporters, as well as validated behavioral tests that can be used to relate injuries to organism-wide consequences and outcomes relevant to humans (Egan et al., 2009). Indeed, zebrafish have long been used as models for traumatic brain and spinal cord injuries, but as in mouse models, most zebrafish models developed to date either penetrate the skull or are technically prohibitive and focus on damage to nervous tissues (Baumgart et al., 2012; Cacialli et al., 2018; McCutcheon et al., 2017; Mokalled et al., 2016).

Here, we leverage the advantages of the zebrafish, in particular the ability to image the meninges through the intact skull of adult zebrafish, to develop a powerful new model for live longitudinal imaging of mCVI and its longer-term sequelae. This new injury model creates small, sterile meningeal cerebrovascular injuries that result in meningeal vascular disruption and meningeal bleeding with no detectable damage to the brain, effectively modeling the minor head wounds and meningeal cerebrovascular trauma that frequently occurs in humans. To accomplish this, we adapted and calibrated a dental sonication tool to induce focal mCVI. Behavioral analysis after injury show these injuries elicit only minor aberrant anxiety responses, reflecting the lack of brain damage we observe. We show that transgenic vascular and immune reporter lines can be used for high-resolution longitudinal live imaging post-mCVI.

## RESULTS

### An adult zebrafish model of sterile dural meningeal cerebrovascular injury

In humans, meningeal cerebrovascular injury (mCVI) often involves damage to the dural venous sinuses (Arbona-Lampaya et al., 2025; Turtzo et al., 2025). Like humans, adult zebrafish have well-developed dural venous sinuses, as readily visualized by confocal imaging of vessels through the translucent skull of *Tg(kdrl:mcherry)^y205^* transgenic (Fujita et al., 2011), melanocyte-free *casper (White et al., 2008)* animals **(Fig. 1A-D)**. The transverse sagittal sinus (TSS) is in the dura above the cerebral ventricles, while the superior sagittal sinus (SSS) runs between the optic tecta, with smaller tributaries above the cerebellum in some individuals **(Fig. 1A-B, Supp. Fig. 1A-E)**. The two perpendicular sinuses meet in a plexus of vessels at the confluence of sinuses (COS). While every fish has the same gross anatomy of dural vascular sinuses, similarly to humans the detailed patterning of the vessels varies between individuals **(Supp. Fig. 1)**.

**Figure 1.**
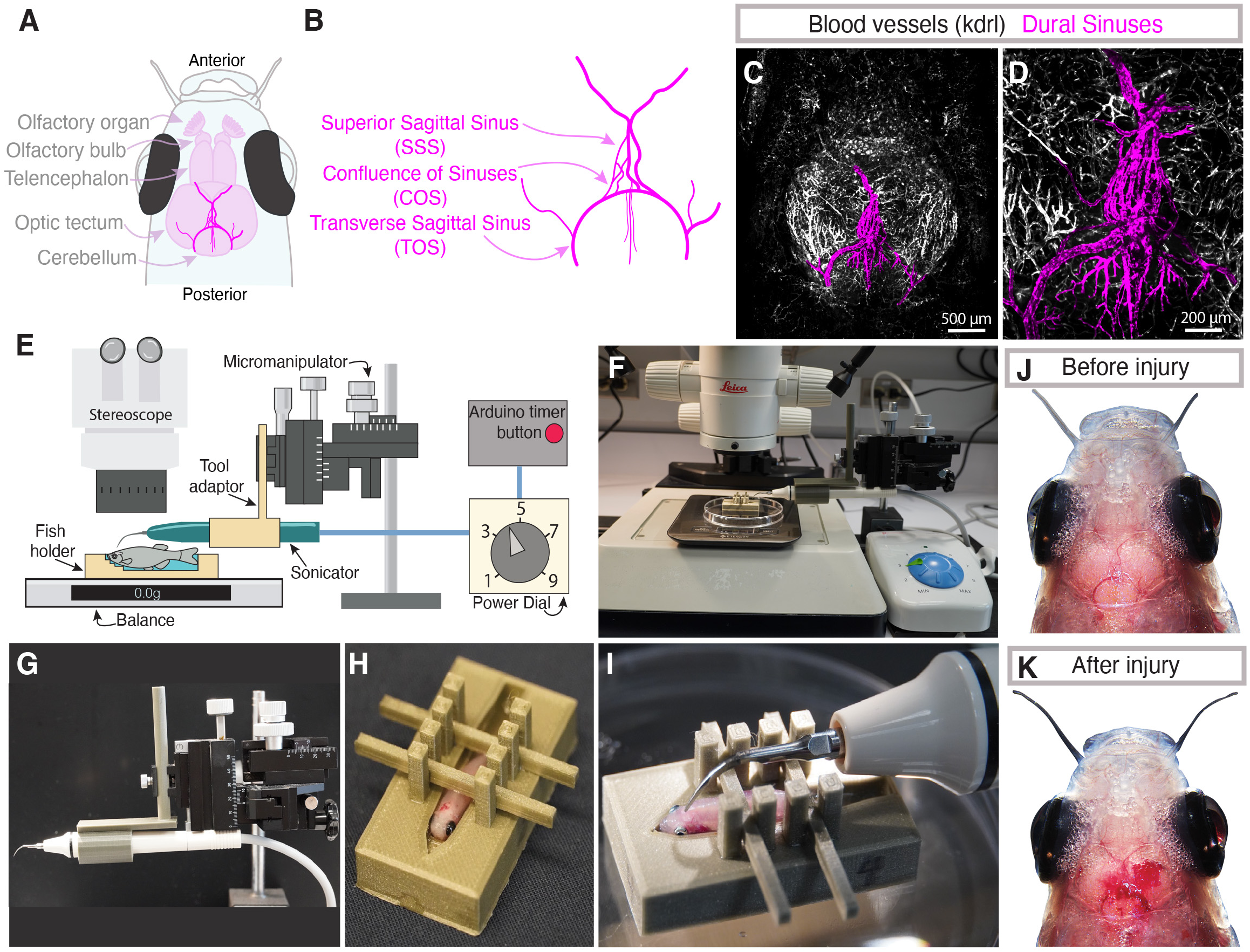
An adult zebrafish model of dural meningeal cerebrovascular injury. **A,B,** Schematic diagrams of the dorsal adult zebrafish brain (A) and the dural vasculature (B), highlighting the presence of the transverse sagittal sinus (TOS) running around the cerebellum, the superior sagittal sinus (SSS) between the two optic tecta, and the confluence of sinuses (COS) where the TOS and SSS meet. **C,D,** Confocal images of the dorsal head of a 6-month-old *Tg(kdrl:mcherry)^y205^*animal, with the dural sinus vasculature pseudocolored in magenta. Panel D shows a magnified view of a portion of the image in panel C. Scale bars are 500μm (C) and 200μm (D). **E,** Schematic diagram of the meningeal cerebrovascular injury (mCVI) apparatus and its composite parts. **F,** Overview image of the mCVI apparatus set up under a stereomicroscope. **G,** Image of the ultrasonic dental scaler attached to a 3D printed adaptor mounted on a micromanipulator. **H,** Image of an adult zebrafish mounted in a 3D printed fish holder with braces to hold the animal in place for the mCVI procedure. **I,** Image of an adult zebrafish with the tip of the dental tool placed positions above the COS for mCVI. **J,K,** Images of the dorsal head of a zebrafish before (J) and after (K) the mCVI procedure, highlighting the highly transparent skull with the brain and major blood vessels clearly visible and the presence of meningeal hemorrhage after mCVI in (K). See **Supp. Movie 1** for videography of a typical mCVI procedure.

We repurposed an ultrasonic dental tool to establish an efficient and inexpensive method to generate localized, reproducible dural mCVI lesions (**Fig. 1E-I, Supp. Files 1-12**). A custom 3D printed plastic adaptor (**Supp. File 12**) is used to attach the probe from a Woodpecker UDS-K Ultrasonic Scaler to a stand-mounted micromanipulator for precisely controlled placement of the probe tip **(Fig. 1E,G,I).** To further facilitate positioning of the probe tip and ensure consistent transduction of dental tool power into head tissue, the adult zebrafish is stabilized by mounting anesthetized animals in custom 3D printed plastic fish holders designed in multiple sizes to accommodate different male and female body forms **(Fig. 1H,I, Supp. Files 1-11)**. The fish holder is placed on a balance scale to measure and control the exact pressure applied by the probe tip on the fish’s head. Three grams of pressure allows for effective lesion generation without breaching the skull at the power levels we employ (see below). A custom-built Arduino controller (**Supp. Fig 2**) is used to control the length of ultrasonic pulses (programmed for 1 second pulses; **Supp. File 14**). The entire assembly is placed under a stereomicroscope for magnified observation of the procedure (**Fig. 1E,F)**. This method permits consistent dental tool placement at the same location and rapid generation of reproducible lesions in multiple adult fish **(Fig. 1I)**. Meningeal hemorrhage is clearly visible after inducing mCVI **(Fig. 1J-K).** The entire procedure (mounting of fish, injury induction, returning of fish to holding tank) can be carried out in approximately 3 minutes **(Supp. Movie 1)**, and as further discussed below animals given localized, mild mCVI recover from anesthesia and survive.

**Figure 2.**
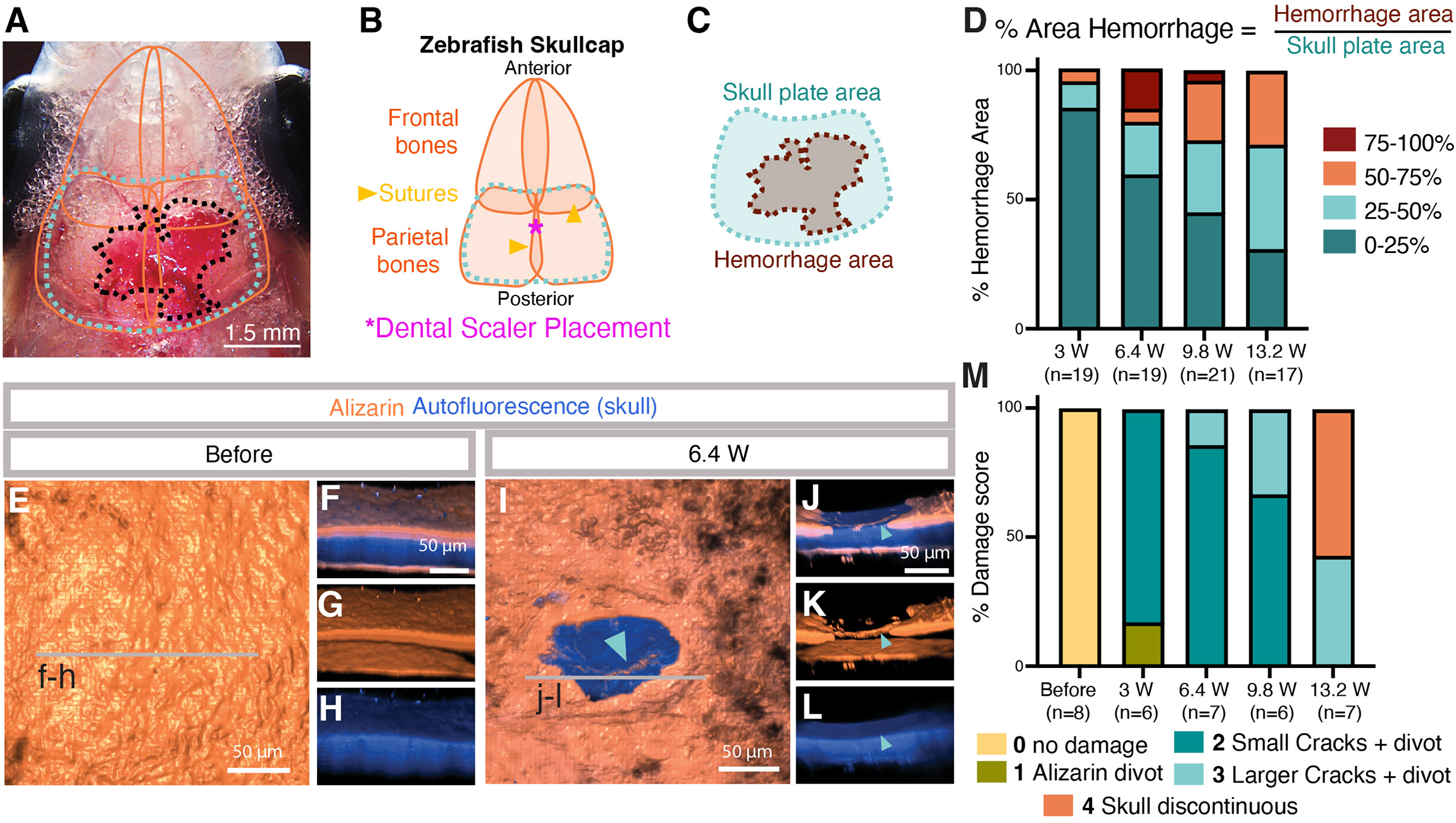
Dural meningeal cerebrovascular injury is scalable and sterile. **A,** Representative image of the dorsal head of a zebrafish with hemorrhage after mCVI (same image as Fig. 1J). The orange lines show the perimeters of the skull bones that lay beneath the skin, the teal dotted line outlines the parietal skull plate area used to quantify hemorrhage area relative to fish size, and the black dotted line indicates the area of hemorrhage (see Fig. B-D for description of measurements). Scale bar = 1.5 mm. **B,** Schematic diagram of the parietal and frontal skull bones. The individual skull bones come together at sutures. The dental scaler tip is positioned on the seam of the two parietal bones over the COS (magenta asterisk). **C,** Schematic diagram of the parietal skull plate and hemorrhage areas used to quantify hemorrhage relative to the size of the animal and its meninges. **D,** Quantification of % hemorrhage area normalized to skull plate area at 4 different dental tool power settings (3, 6.4, 9.8, or 13.2 Watts). The % hemorrhage area was categorized as either 0-25%, 25-50%, 50-75%, or 75-100% of the skull plate area. Numbers of animals measured at each power level (n) are listed below the graph. **E-L,** Confocal imaging of alizarin red-stained skull surface bone (orange) and deeper autofluorescent skull bone (blue) on the dorsal skull of a wild type adult zebrafish before (E-H) or immediately after (I-L) 6.4 W mCVI. Images shown in panels are dorsal views 3D rendered using to show a “solid” view of skull surface (E,I) or lateral views of a portion of the same confocal image stacks beginning from the lines on panels E and H, showing alizarin + autofluorescence (F,J), alizarin alone (G,K), or autofluorescence alone (H,L). Scale bars = 50 µm. See **Supp. Movie 2** for 3D renderings of representative skull image data. **M,** Quantification of skull damage after mCVI at 4 different dental tool power settings (3, 6.4, 9.8, or 13.2 Watts). The scoring criteria are explained in detail in Supp. Fig. 2 and in the accompanying Methods and Results sections. Numbers of animals measured at each power level (n) are listed below the graph.

### The ultrasonic dental scaler generates scalable meningeal injuries in adult zebrafish

We optimized dental tool power to consistently generate localized, “sterile” injuries restricted to the meninges that do not breach the skull. To quantify hemorrhage, we normalized the visible blood (red) in stereoscope images to overall fish size by calculating the percent hemorrhage area beneath the parietal skull plates **(Fig. 2A-C, Supp. Fig. 3A-D)**. The dental tool was positioned above the COS near the intersection of the parietal plates along the skull suture **(Fig. 2B)**. We carried out blinded scoring of hemorrhage at four different tool power (in Watts) levels (3 W, 6.4 W, 9.8 W, or 13.2 W treatments for 1 second each) and found that more power generally increased the likelihood of a large hemorrhage **(Fig. 2D)**. We also assessed skull integrity post-injury by using alizarin red (calcium) live staining and the natural autofluorescence of the skull to visualize skull bone (Castranova et al., 2021) at the injury site **(Fig. 2E-L, Supp. Fig. 3)**. As we have shown previously (Castranova et al., 2021), brief alizarin red treatment of live animals only stains the outer edges of the skull bone, while the entire thickness of the skull bone is visualized by autofluorescence, allowing for detailed assessment of skull damage and integrity (**Fig. 2F**). A blinded scoring system was used to evaluate skull damage **(Fig. 2M, Supp. Fig. 3F-Z)**. In short, the score is 0 if the alizarin and autofluorescence are entirely intact and there is no damage, 1 when there is a small divot in the outer alizarin stained layer but the full skull autofluorescence is intact, 2 when there is an when there is an alizarin divot as well as small very superficial cracks evident in the inner blue autofluorescent layer, 3 when cracks are larger, penetrating halfway into the full thickness of the skull but not entirely penetrating the skull, and 4 when the fluorescent signal is discontinuous and the full thickness of the skull is broken **(Fig. 2M, Supp. Fig. 3F-Z)**. At 6.4 W there are only minor divots and hairline cracks in the external surface of the alizarin red skull stain, but the internal autofluorescent layers remain intact **(Fig. 2I-M, Supp. Movie 2)**. At 13.2 W cracks penetrate through the skull approximately half of the time **(Fig. 2M)**. Because we wanted a non-penetrative, sterile mCVI model, for all subsequent mCVI procedures described in this study we used 1 second treatment at 6.4 W power with 3 grams of pressure to create mCVIs in 8-14 month old adult animals.

**Figure 3.**
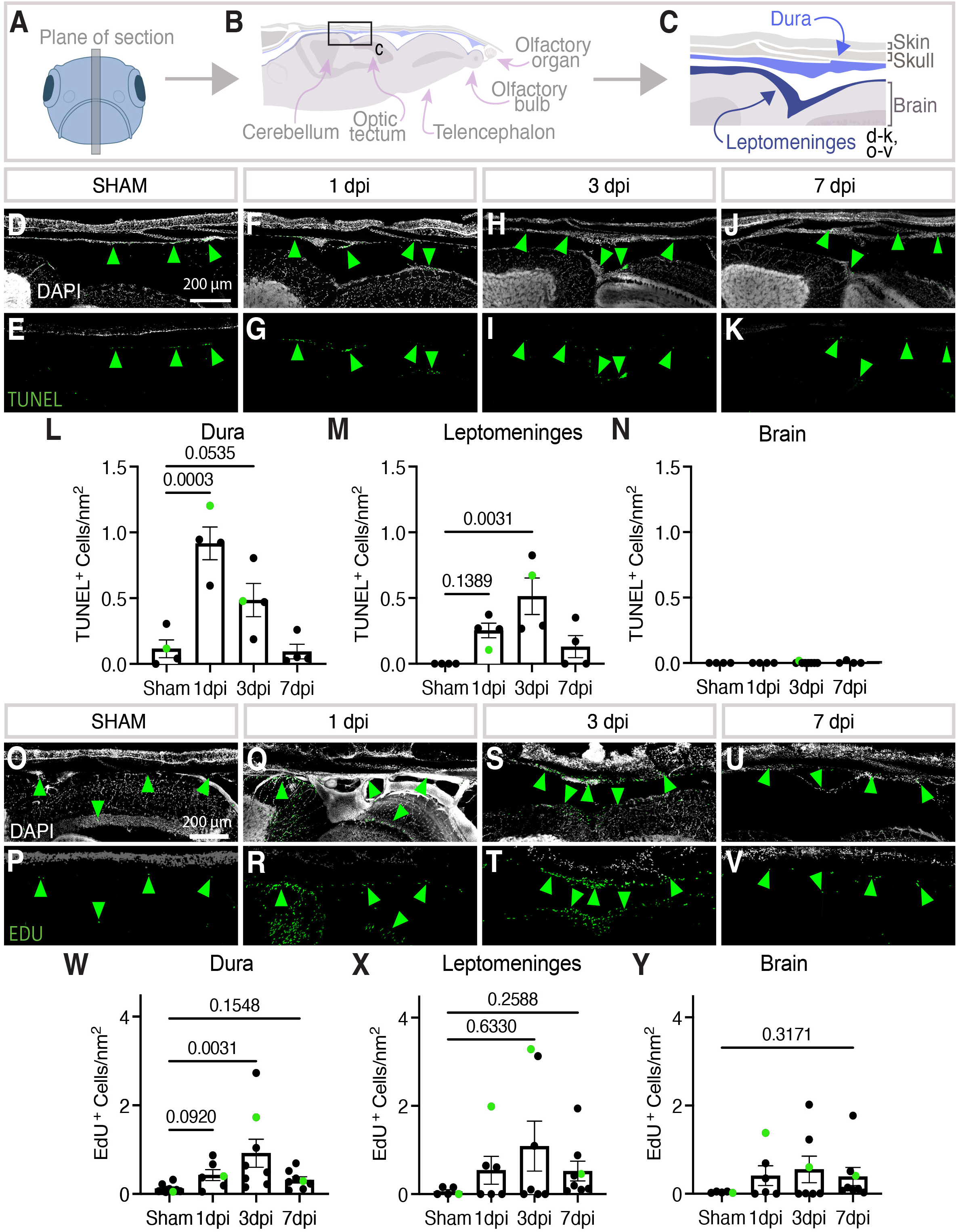
Cell death and cell proliferation after meningeal CVI. **A-C**, Schematic diagrams showing (A) the plane of vibratome sectioning used to collect the data in panels D-Y from adult zebrafish, (B) depiction and annotation of brain and dorsal skull tissues present in vibratome sections, (C) magnified view of the dorsal head from the schematic in panel B with dura and leptomeninges layers highlighted in blue, corresponding roughly to the fields of view in panels D-K and O-V. **D-K,** Confocal images of DAPI- and TUNEL-stained vibratome sections prepared as noted in panels A-C, with DAPI-stained nuclei in white (D,F,H,J) and TUNEL staining in green (D-K). Sections imaged are from sibling adult animals that were either uninjured (D,E) or 1 (F,G), 3 (H,I), or 7 (J,K) days post-mCVI (dpi). Scale bars are 200μm. **L-N,** Quantification of the number of TUNEL-positive cells per mm^2^ of section in the dura (L), leptomeninges (M), and adjacent brain parenchyma (N). Statistics are ordinary one-way ANOVA with Dunnett’s multiple comparisons *post hoc* test. **O-V,** Fluorescence microscopy of DAPI- and 5-ethynyl-2′-deoxyuridine (EDU)-stained vibratome sections prepared as noted in panels A-C, with DAPI-stained nuclei in white (O,Q,S,U) and EDU staining in green (O-V). Sections imaged are from sibling adult animals that were either uninjured (O,P) or 1 (Q,R), 3 (S,T), or 7 (U,V) days post-mCVI (dpi). Scale bars are 200μm. **W-Y,** Quantification of the number of EDU-positive cells per mm^2^ of section in the dura (W), leptomeninges (X), and adjacent brain parenchyma (Y). Statistics are non-parametric with Kruskal Wallis with Dunn’s multiple comparisons *post hoc* test.

### Intracranial damage is largely restricted to the meninges

We carried out additional characterization of our model to determine whether damage is localized to the meninges or if the brain parenchyma is also being injured. Vibratome sections were collected from the midline anterior-posterior axis of fixed injured fish heads **(Fig. 3A-C).** These did not reveal obvious swelling of tissues adjacent to meningeal hematomas, but to assess the level of cell death we used TUNEL staining to measure the number of apoptotic cells in the dura, leptomeninges, and the adjacent cerebellar brain tissue in sham-treated or 1-, 3-, 7-day post injury (dpi) animals **(Fig. 3D-N)**. Interestingly, apoptosis peaks rapidly at 1 dpi in the dura, but peaks at 3 dpi in the leptomeninges **(Fig. 3L,M)**. Importantly, virtually no cell death was observed in the adjacent brain parenchyma **(Fig. 3N)**. We also evaluated cell proliferation as a proxy for healing **(Fig. 3O-Y)**. We carried out intraperitoneal injection of 5-ethynyl-2′-deoxyuridine (EdU) 24 hours prior to euthanasia and sacrificed age-matched siblings on the same day to control for baseline proliferation in the adult zebrafish brain. Staining of longitudinal sections reveal that proliferation peaks at 3 dpi in the dura and leptomeninges **(Fig. 3W,X)**. The number of EdU^+^ cells was also increased in the brain parenchyma at 3 dpi, suggesting that despite the lack of cell death in the parenchyma there may be some indirect or direct effects on cells in the parenchyma **(Fig. 3Y)**. Given that initial cell death is concentrated in the dura, we believe our model primarily damages the dural meninges.

### Aberrant anxiety behavior is associated with mild meningeal cerebrovascular injury

Mammals exhibit anxiety behaviors after head injuries (Delmonico et al., 2022; Osborn et al., 2017; Popovitz et al., 2019; Scholten et al., 2016). To assess whether zebrafish subjected to mCVI develop these behaviors, we used the novel diving tank test (Anwer et al., 2021; Egan et al., 2009), a commonly used assay for measuring anxiety behavior in adult zebrafish **(Fig. 4A, Supp. Movie 3)**. In this test, more anxious zebrafish should linger in the bottom of the tank longer than less anxious fish. Anxiety read-outs include both latency to enter the top half of the tank (defined as the time from when the fish settles at the bottom to when it first crosses into the top) and percentage of total time spent in the top half of the tank (Cachat et al., 2010; Egan et al., 2009). For each fish, anxiety was measured longitudinally at baseline and at 1, 3, 7, 13, and 20 days post-mCVI or sham treatment. As a positive control for anxiety responses fish were chased for 30 seconds with a 3D-printed plastic toy predator (**Supp. File 13**)(Tran et al., 2014) before the novel diving tank test 22 days after either mCVI or sham treatment. Representative swim traces show that mCVI treated animals spent more time moving slowly and lingering at the bottom of the tank, while sham-treated animals maintained consistent movement patterns across longitudinal testing **(Fig. 4B-F)**. Latency to the top of tank was slightly faster after CVI compared to sham-treated animals in the first week, although only significant at 7 dpi **(Fig. 4G)**, and this effect was reduced over subsequent time. Mean velocity and therefore distance swum remained relatively constant, suggesting that CVI did not impair locomotion **(Fig. 4H)**. Strikingly, the percent of time spent in the top half of the tank was significantly less for mCVI treated animals compared to sham treated **(Fig. 4I)**. Together these results suggest there is a very small but measurable increase in anxiety-associated behavior after modest dural mCVI.

**Figure 4.**
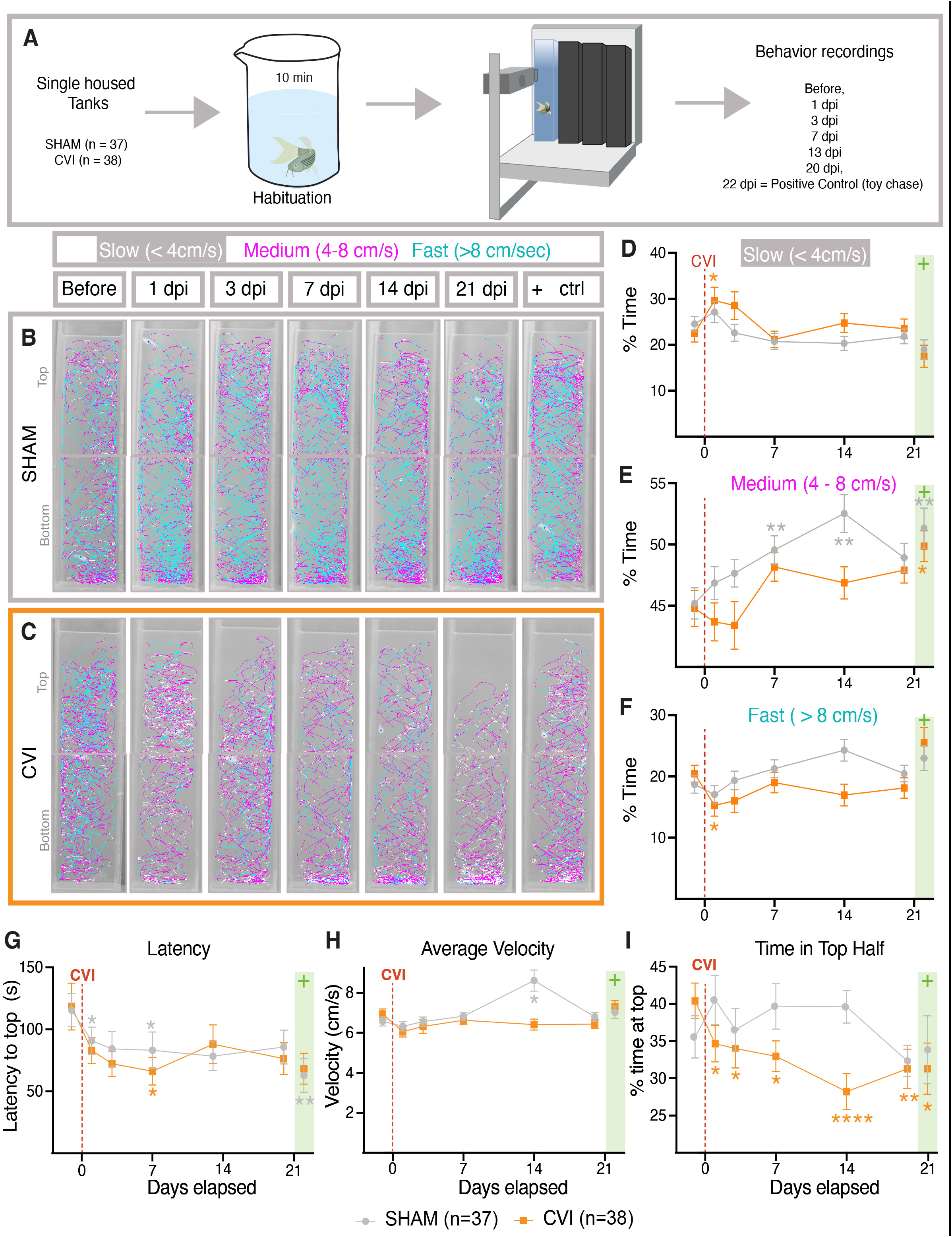
Anxiety responses after meningeal cerebrovascular injury. **A,** Schematic diagram of the diving tank test used to measure anxiety where individually housed animals are habituated in a beaker for 10 min before being introduced to the diving tank and having their movements recorded for 10 min. All animals were measured at initial baseline and at 1, 3, 7, 13, and 20 days after either sham or mCVI treatment. On day 22 post-treatment, fish were chased with a toy predator for 30 seconds before being netted for habituation and measurement. See **Supp. Movie 3** for representative live images of behavior testing. **B,C,** Tracks from representative sham (B) and mCVI (C) treated animals showing the locations of the animals and their approximate speeds (white, < 4 cm/s, pink, 4-8 cm/s, and teal, > 8 cm/s). **D-I,** Quantification of behavioral data for sibling animals at initial baseline and at 1, 3, 7, 13, and 20 days after either sham or mCVI treatment. Graphs show (D) percent of time going at slow speed (<4 cm/s), (E) percent of time going at medium speed (4-8 cm/s), (F) percent of time going at fast speed (>8 cm/s), (G) latency (time in seconds) to cross to the top half of the tank, (H) average velocity (cm/second), (I) percent of time spent in the top half of the diving tank. Sham or mCVI treatment on day 0 is noted with a red dashed line. The toy chase on day 22 is noted with green box. For sham group n = 37 and fpr mCVI n = 38 across 2 independent experimental runs. Statistics are a two-way ANOVA with Dunnett test for multiple comparisons. * p < 0.05, ** p < 0.01, **** p < 0.0001.

### Live imaging of the vasculature after meningeal cerebrovascular injury

Our model and the availability of transgenic reporter lines for endothelial or blood cells permits longitudinal live confocal imaging of dural hemorrhage and dural vascular damage, regrowth, and remodeling in the same animal after mCVI. Imaging of vibratome sectioned *Tg(kdrl:egfp)^la116^*, *Tg(gata1:dsred)^sd2^* double transgenic adults confirms extensive hemorrhage after mCVI. Unlike sham-treated animals in which gata1:dsred-positive blood cells are only found within kdrl:egfp-positive blood vessels **(Fig. 5A-C)**, large numbers of extravascular blood cells are clearly visible in the vicinity of the injury immediately after mCVI **(Fig. 5D-F)**. By 1 dpi there are already many fewer intact blood cells in the meninges, although a substantial amount of debris from these cells appears to remain **(Fig. 5G-I)**. Blood cells can also be fluorescently “tagged” and live-imaged concurrently with vessels through the translucent skull by intravenous injection of Hoechst nuclear dye into a *Tg(kdrl:egfp)^la116^* transgenic, *casper* adult animal, confirming extensive extravasation of blood from damaged dural vessels immediately after mCVI **(Fig. 5J-L, Supp. Movie 4)**.

**Figure 5.**
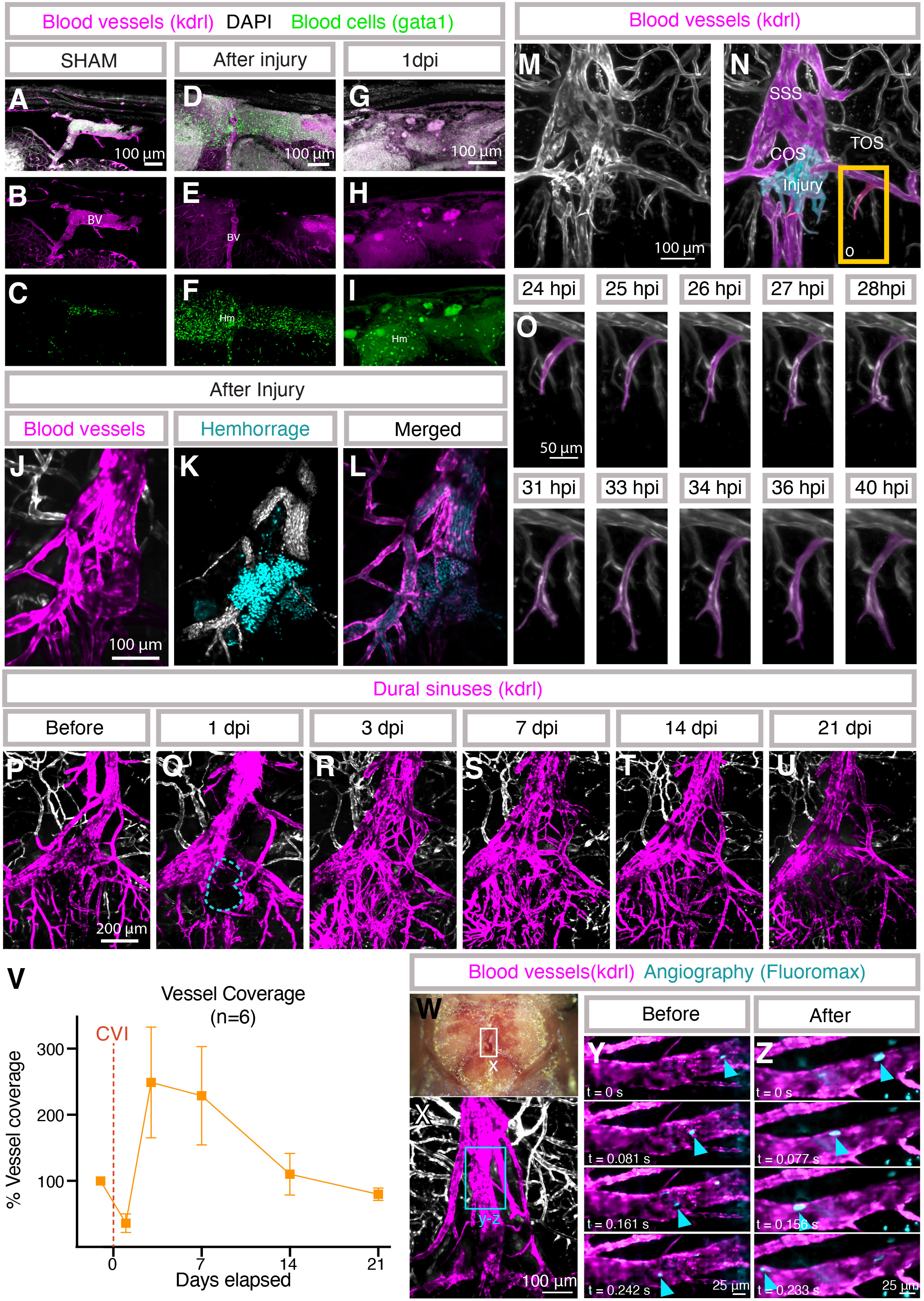

Time-lapse imaging of an intubated (Castranova et al., 2022) *Tg(kdrl:mcherry)^y205^*transgenic *casper* adult animal from 24-40 hours post-injury (hpi) reveals active angiogenesis of a transverse sagittal sinus (TOS) tributary vessel near the site of injury, in contrast to the quiescent dural vasculature of an uninjured animal **(Fig. 5M-O, Supp. Movie 5)**. Longitudinal assessment of dural vessels can also be carried out over the course of weeks or potentially even months by repeated intubation and imaging of the same animal. We imaged dural blood vessels in a *Tg(kdrl:mcherry)^y205^* transgenic *casper* adult animal just before injury and at 1, 3, 7, 14, and 21 days post-mCVI. The avascular area is revascularized as rapidly as 3 dpi **(Fig. 5P-V)**. By 7dpi, there is an overgrowth of vessels that resolves to close to baseline vascular density by 14 dpi, although some of the vessels are more tortuous compared to the aligned architecture of the dural venous sinus before injury **(Fig. 5S, T)**. By 21 dpi the vascular area has returned to pre-injury vascularity with a similar level of plexus complexity **(Fig. 5U-V)**.

Despite the extensive hemorrhage and evident dural vascular rupture in our model, adult zebrafish all survive injury under our “standard conditions” (1 second treatment at 6.4 W power with 3 grams of ultrasonic probe tip pressure), suggesting local blood supply to the dura is largely preserved. To investigate this, we live-imaged blood flow in the superior sagittal sinus (SSS) of an animal immediately (10 minutes) after mCVI (**Fig. 5W-Z; Supp. Movie 6**). The SSS is just anterior to the site of mCVI injury at the confluence of sinuses (COS; see **Fig. 1A,B** for anatomical reference). After intravascular injection of 20 µm FluoroMax beads we see that robust blood flow is still present in the SSS downstream from the mCVI site, suggesting that despite extensive hemorrhage dural vessels are able to maintain or rapidly restore blood supply.

### Live imaging of immune cells after meningeal cerebrovascular injury

Much of the pathology of human mCVI is the result of the inflammatory sequelae that follow the initial injury-induced vascular rupture and hemorrhage (Llull et al., 2025; Melmed et al., 2021). Understanding how immune cells are recruited and how they function at the site of injury to facilitate or moderate this inflammatory response is of vital importance but difficult to study in living mammalian models. As a first test to determine whether our model could be used to carry out longitudinal live imaging of immune cell dynamics following mCVI, we imaged neutrophils in *Tg(lyz:dsred2)^nz50^* transgenic (Hall et al., 2007) animals after dural mCVI. Neutrophils are only sparsely present in the major dural vasculature of uninjured zebrafish, as in mammals (Smyth et al., 2024; Venero Galanternik et al., 2025), and they are similarly quiescent **(Fig. 6A-B)**. However, within a few hours after mCVI lyz:dsred-positive neutrophils rapidly accumulate within and near the wounded vasculature **(Fig. 6B-G, Supp. Movie 7)**. By 3 hours post-injury (hpi) numerous neutrophils can be seen rolling through lumens of nearby blood vessels **(Fig. 6C, Supp. Movie 7)**. By 10 hpi neutrophils have “swarmed” the site of CVI in the meninges, presumably extravasating from circulation **(Fig. 6F)**. Measurement of neutrophil numbers just before injury and at 1, 3, 7, 14, and 21 days post-mCVI (dpi) shows that neutrophils rapidly peak at 1 dpi, resolving to near-baseline levels by 21 dpi (**Fig. 6H-M,T)**. The dynamics of individual neutrophils can also be measured longitudinally, including neutrophil transit speed through the injured dural venous sinuses. Live imaging and tracking of neutrophils in the dural venous sinuses of intubated *Tg(lyz:DsRed2)^nz50^*transgenic adults just before injury and at 1, 3, 7, 14, and 21 dpi reveals observed a substantial decrease in neutrophil speed through the SSS at 1 dpi that is largely reversed by 3 dpi, and that has returned to steady state by 21dpi **(Fig. 6N-S,U, Supp. Movie 8)**. A transient decrease in the speed of neutrophils transiting vessels after injury is a well-documented behavior of mammalian neutrophils indicative of increased interactions between blood vessels and leukocytes.

**Figure 6.**
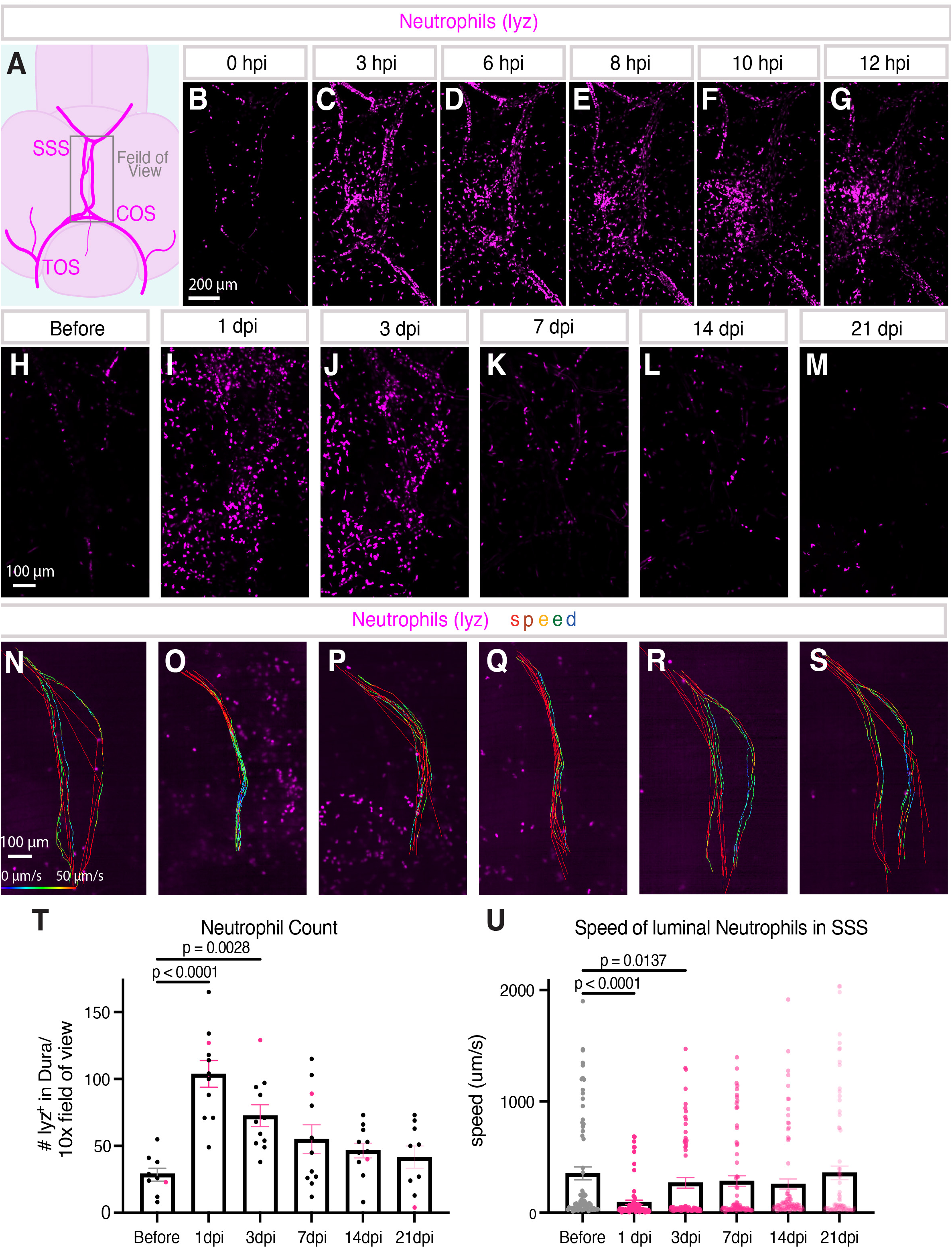
Vascular responses to meningeal cerebrovascular injury. **A-I,** Confocal images of DAPI-stained vibratome sections from *Tg(kdrl:egfp)^la116^, Tg(gata1:dsred^sd2^)* double transgenic adult zebrafish prepared as noted in **Fig. A-C**, with DAPI-stained nuclei in white (A,D,G), kdrl-positive blood vessels in green (A,B,D,E,G,H), and gata1-positive blood cells in magenta (A,C,D,F,G,I). Sections imaged are from sibling animals that were either uninjured (A-C), immediately post-mCVI (D-F) or 1 day post-mCVI (G-I). BV = blood vessels, Hm = hemorrhage. Scale bars = 100 μm. **J-L,** Confocal image of hemorrhage at the confluence of sinuses (COS) collected through the skull of a living Hoechst dye-injected *Tg(kdrl:mcherry)^y205^* transgenic adult immediately after mCVI. Kdrl-positive COS vessels are in pseudocolored in magenta in panel J, circulating blood cells are in white and extravasated blood cells in teal in panel K, and blood vessels magenta and blood cells teal in panel L. Scale bars = 100μm. See **Supp. Movie 4** for a live movie of circulation through the same vessels. **M-O,** Time lapse-confocal images of regrowing dural vessels collected through the skull of a living *Tg(kdrl:mcherry)^y205^*transgenic adult 24 hours after mCVI. Panels M and N show an overview image of the dural vessels in the area of injury at the start of time-lapse imaging, with dural vessels in magenta and the site of injury in teal in panel N. Panel O shows selected frames from time lapse images of a regrowing vessel in the boxed region in panel N. Scale bars = 100 µm (M,N) and 50 µm (O), abbreviations in panel N are as noted in Fig. 1B. See **Supp. Movie 5** for the full time lapse series. **P-U,** Live longitudinal time-series confocal images of dural vessel (pseudocolored magenta) regrowth and remodeling, collected through the skull of the same living *Tg(kdrl:mcherry)^y205^* transgenic adult immediately before (P) and at 1 (Q), 3 (R), 7 (S), 14 (T), and 21 (U) days post-mCVI (dpi). **V,** Quantification of vascular coverage in 6 separate *Tg(kdrl:mcherry)^y205^* transgenic adults (including the animal shown in panels P-U) immediately before and at five additional time points post-mCVI. Coverage is shown as percentage of the initial vascular coverage before mCVI, measured from normalized fluorescence of *kdrl:mcherry* in a region of interest drawn around the avascular area observed in each animal at 1 dpi. **W-Z,** Real-time confocal imaging of blood flow though the superior sagittal sinus (SSS) before and immediately after mCVI in the nearby confluence of sinuses (COS), collected through the skull of a living *Tg(kdrl:mcherry)^y205^* transgenic adult intravascularly injected with FluoroMax beads. (W) Stereoscope image of the head of the animal live imaged in panels X-Z, with a box around the region imaged in panel X. (X) Confocal image of dural vessels (pseudocolored magenta). The region including a portion of the superior sagittal sinus (SSS) shown in panels Y and Z is noted with a box. Scale bar = 100μm. (Y,Z) Sequential frames from live confocal imaging of a segment of the SSS in the same animal before (Y) and immediately after (Z) mCVI, showing robust flow post-mCVI. Blue arrowheads show movement of a fluorescent bead. Scale bar = 25μm. See **Supp. Movie 6** for the full live image series.

## DISCUSSION

Here we present a novel model of meningeal cerebrovascular injury (mCVI) in adult zebrafish generated with an ultrasonic dental scaler. This approach produces focal, sterile vascular injuries in the meninges without breaching the skull or causing cell death in the brain. Although the brain parenchyma remains intact, behavioral assays using the novel tank test revealed a modest increase in anxiety-like behavior after mCVI, suggesting functional consequences of meningeal injury alone. The translucent adult zebrafish skull permits real-time, high-resolution optical imaging of vascular leakage and repair in mammalian-like meninges without invasive surgery. Using transgenic reporter lines and intubation methods, we observed reproducible vascular overgrowth followed by pruning over subsequent days to weeks, as well as rapid neutrophil recruitment through the vasculature. Together, this model provides a tractable and versatile system for dissecting the vascular–immune interactions and inflammatory responses that shape recovery after mCVI.

The meninges are increasingly recognized as active immune–vascular hubs rather than passive coverings of the brain. In zebrafish, the meninges display mammalian-like anatomy, with a highly vascularized dura attached to the skull and underlying leptomeningeal layers (Venero Galanternik et al., 2025) Like mammals, zebrafish possess dural venous sinuses that drain blood from the meninges and superficial brain, and we observed inter-individual variability in this network similar to that seen in humans and mice (Brockmann et al., 2012; Jacob et al., 2022; Mecheri et al., 2018; Shapiro et al., 2021) (**Supp. Fig. 1**). These venous structures are closely associated with immune compartments and have been implicated in disorders including infection and meningiomas (Caroli et al., 2006; Coles et al., 2017; Fitzpatrick et al., 2020; Fitzpatrick et al., 2024). The zebrafish meninges also contain a full repertoire of innate and adaptive immune cells, mirroring human and mouse systems (Rustenhoven et al., 2021; Schafflick et al., 2021; Venero Galanternik et al., 2025). This conservation highlights the zebrafish as a powerful model for studying meningeal vascular and immune dynamics in the context of health and disease.

We sought to establish a reproducible model of mild meningeal cerebrovascular injury (mCVI) with scalable control of injury severity. Using an ultrasonic dental scaler with Arduino-timed power control (**Fig. 1**), we found that increasing power levels produced progressively larger hemorrhages (**Fig. 2D**). At lower settings, this approach generated a closed-skull, “sterile” meningeal wound, while higher levels occasionally breached skull (**Fig. 2E–M**). Given that human brain injuries can range from mild and survivable to severe and fatal (Maas et al., 2008; Siebold et al., 2018), we tuned the model to reproducibly induce a mild mCVI in which all animals survived, brain parenchyma remained intact, and behavioral changes were modest (**Fig. 3N, 5D–I**). Notably, even large hematomas had limited vascular damage, showing rapid sealing of dural sinus injury and maintained blood flow by angiography (**Fig. 5J–L, Supp. Movies 4,6**). This establishes a tractable and physiologically relevant model for studying vascular–immune interactions following mild mCVI.

Zebrafish are already established as neuroimmune models for brain injury, including stroke, ischemia, and spinal cord trauma, owing to their regenerative capacity and amenability to high-throughput testing (Cacialli, 2021; Cho et al., 2020; Crilly et al., 2022; de Medeiros Borges et al., 2023; Gan et al., 2020; Kishimoto et al., 2012; Kizil et al., 2012; Murashova and Dyachuk, 2025; Skaggs et al., 2014; Zhu et al., 2020; Zulazmi et al., 2021). Prior zebrafish models of traumatic brain injury, including focused ultrasonic injuries that are technically complex, have been directed at studying the brain parenchyma rather than the meninges (Cho et al., 2020; McCutcheon et al., 2017). Similarly, most vertebrate brain injury models have emphasized neuronal regeneration or inflammation, with limited focus on the impact of vascular damage (Crilly et al., 2022; Crilly et al., 2018; Gan et al., 2020; Marz et al., 2011). Yet hemorrhage in and around the brain is known to drive inflammation, metabolic stress, neuronal death, and poor outcomes when unresolved (Chen et al., 2022a; Livingston et al., 2017; Russo et al., 2018). Our model directly addresses this gap by creating a precise meningeal vascular injury that disrupts meningeal vessels without killing underlying neurons (**Fig. 3N**). Despite the brain remaining intact, we observed proliferative responses in adjacent tissue (**Fig. 3Y**), consistent with the high regenerative potential of zebrafish (Becker and Becker, 2022; Kishimoto et al., 2012; Skaggs et al., 2014; Zambusi and Ninkovic, 2020). Unlike mammals, where neuronal precursor cells proliferate after injury but rarely integrate long-term (Arvidsson et al., 2002; Kojima et al., 2010), zebrafish progenitors migrate to injured networks, integrate, and differentiate (Kishimoto et al., 2012; Skaggs et al., 2014), making them uniquely suited to study how vascular injury and immune responses influence neural repair. Our approach therefore provides the first tractable zebrafish system to specifically model meningeal vascular injury and its associated immune responses.

Because even mild traumatic brain injuries in humans and rodents are associated with anxiety (Delmonico et al., 2022; Osborn et al., 2017; Popovitz et al., 2019; Scholten et al., 2016), we tested whether zebrafish exhibit similar behaviors after mCVI. We used the novel diving tank assay **(Supp. Movie 3)** (Cachat et al., 2010; Egan et al., 2009) with tall, narrow infrared tanks (52 cm × 7 × 10.5 cm) previously shown to increase sensitivity in detecting anxiety-like responses (Anwer et al., 2021). While reduced light at the bottom introduced a scototaxis variable (Maximino et al., 2010), light–dark transitions are also used as an anxiety measure, suggesting this did not confound interpretation. The total distance swum and average velocity did not differ between sham and CVI fish (**Fig. 4H**), confirming locomotor function was intact. After mCVI, fish spend less time swimming at high speeds and more time swimming at slower speeds needed to turn more frequently, resembling the “meander” behavior described in other mild brain injury models (McCutcheon et al., 2017) (**Fig. 4B–F**). Latency to enter the top half of the tank was significantly shorter in CVI fish at 7 dpi (**Fig. 4G**). Both sham and CVI groups showed reduced latency over time, consistent with habituation, but the stronger effect in CVI fish suggests that mild meningeal injury amplifies behavioral adaptation beyond habituation alone (**Fig. 4G**). Despite this, CVI-treated fish spent significantly less time in the top half compared to shams, preferring the dark bottom zone and exhibiting less exploratory behavior (**Fig. 4I**), a classic indicator of increased anxiety in fish (Egan et al., 2009; Maximino et al., 2010). Thus, although modest, anxiety-like behaviors increased in zebrafish following mCVI. Importantly, these findings show that isolated meningeal vascular injury, even in the absence of overt brain damage, is sufficient to produce measurable behavioral changes. This establishes a functional link between meningeal CVI and altered behavior, paralleling the anxiety phenotypes observed in humans and rodents after mild TBI, and providing a sensitive readout for the downstream impact of vascular–immune interactions in our model.

Studying vascular–immune interactions after cerebrovascular injury (CVI) has been technically challenging: larval zebrafish models are confounded by ongoing development and neutrophil migration through interstitial tissues, while live imaging in adult mice requires skull thinning that induces inflammation. Larval zebrafish have been instrumental for defining neutrophil biology in myriad wound model, particularly their rapid migration to wounds, egress, and roles in modulating macrophage phenotypes (Chen et al., 2022b; de Sena-Tomas et al., 2024; Mathias et al., 2006; Michael et al., 2025; Michael and de Oliveira, 2023; Tsarouchas et al., 2018). To overcome the limitations of larval and mammalian systems, we applied adult zebrafish intubation methods that permit non-invasive, longitudinal imaging of the meninges (Castranova et al., 2022; Castranova et al., 2021; Greenspan et al., 2024). Immediately after mCVI, we observed neutrophils entering the meninges through the dural vasculature and migrating into injured tissues (**Fig. 6A–F; Supp. Movies 7**). We tracked neutrophil infiltration immediately after injury, observing migration through the dural vasculature and into the meningeal tissues **(Fig. 6A-F; Supp. Movies 7)**. Furthermore, because the fish remain unharmed during live imaging, we can perform longitudinal time-lapse imaging to analyze vascular–immune interactions, such as neutrophil movement through the dural sinuses, which was slower on average during peak infiltration **(Fig. 6O-V; Supp. Movies 8)**. Because the skull remains intact, these observations are free from surgically induced inflammation and can be repeated over time in the same animal. Thus, the adult zebrafish mCVI model uniquely enables direct, high-resolution imaging of vascular–immune dynamics in vivo, providing insights not achievable in larval zebrafish or mammalian systems.

In conclusion, we have established a reliable method to generate focal, sterile mCVI in adult zebrafish using an ultrasonic dental scaler. By leveraging the translucent skull, numerous cell type–specific transgenic reporter lines, and recently developed intubation methods, we show that noninvasive, high-resolution imaging of vascular regrowth, remodeling, and immune cell recruitment can be performed repeatedly in the same animal after mild mCVI. Our approach provides a powerful platform for longitudinal analysis of vascular–immune dynamics after mCVI and will enable new insights into the inflammatory mechanisms that shape recovery.

## MATERIALS AND METHODS

### Animal husbandry and strains

Fish were housed in a large zebrafish-dedicated recirculating aquaculture facility (four separate 22,000-liter systems) in 6-liter and 1.8-liter tanks. Fry were fed rotifers, and adults were fed Gemma Micro 300 (Skretting) once per day. Fish were on a typical 14-hour light, 10-hour dark light cycle. Water quality parameters were routinely measured, and appropriate measures were taken to maintain water quality and temperature (80°C) stability (water quality data available upon request). All fish for imaging and behavior were in a *Casper* – *roy*, *nacre* double mutant (White et al., 2008) – genetic background to increase clarity for microscopy by eliminating melanocyte and iridophore respectively. The following transgenic fish lines were used for this study: *Tg(kdrl:mcherry)^y205^*(Fujita et al., 2011), *Tg(kdrl:egfp)^la116^* (Jin et al., 2005)*, Tg(lyz:DsRed2)^nz50^*(Hall et al., 2007), *Tg(mpeg1:EGFP)^gl22^* (Ellett et al., 2011), *Tg(gata1:DsRed)^sd2^* (Traver et al., 2003). While not shown in the manuscript some of these animals had either *Tg(lyve1:DsRed2)^nz101^*(Okuda et al., 2012), *Tg(mrc1a:eGFP*)*^y251^* (Jung et al., 2017) in the genetic background. Transgenic zebrafish were generated on myriad wild-type backgrounds but are maintained in house by out-crossing to the EK wild-type zebrafish. This study was performed in an American Association for Accreditation of Laboratory Animal Care (AAALAC)-accredited facility under an active research project overseen by the National Institute of Child Health and Human Development Animal Care and Use Committee (NICHD ACUC), Animal Study Proposal # 21-015 and # 25-015.

### Ultrasonic Scaler Injury Model

To create meningeal CVI, a Woodpecker UDS-K Ultrasonic Scaler was repurposed. In short, the Woodpecker UDS-K Ultrasonic Scaler’s foot pedal was removed and the wires were instead connected to a custom-designed controller (**Fig. 1E**) consisting of a breadboard (ElectroCookie Solderable Mini Breadboard PCB) with a relay module (AEDIKO Relay Module DC 12V), a button (Adafruit Stemma Wired Tacticle Push Button pack 4431), and an Arduino (Arduino UNO R4 Wifi and Starter Kit K000007) (**Supp. Fig. 2**). The Arduino was programmed to activate and relay power to the scaler for 1 second after pressing the button (**Supp. File 14**). All electronic components were placed inside a repurposed container for their protection. A PD1 scaler tip was used for all injuries, and injuries were performed under a Leica (LEICA DFC7000 T) stereomicroscope for accuracy of placement of the scaler on the zebrafish head.

Designs for 3D-printed fish holders, holding braces, and micromanipulator adaptor are provided (see Supplemental Materials). The printed fish holder was designed with a reservoir to hold tricaine system water (0.04%) to keep the fish anesthetized and gills wet, and with a chin rest to minimize head movement during the mCVI procedure (**Supp. Files 1-10**). Anchoring rods were printed to secure fish to the platform of the fish holder and prevent sliding during injury (**Supp. File 11**). The ultrasonic scaler was attached to a printed micromanipulator adaptor for spatial control (**Supp. File 12**). 3D printed items were designed with SketchUp 3D software (SketchUp for Web, 2025) and printed with polylactic acid on an Original Prusa i3 MK3S+ 3D printer (Prusa Research). Models can also be printed by a commercial supplier using the .stl files that we provide (**Supp. Files 1-13**).

Adult zebrafish aged 6-8 months were anesthetized in 126 mg/L (1x) Tricaine (Tricaine S, version 121718, Syndel; buffered to pH 7 with 1 M Tris-pH9) until they did not react to a fin pinch. The fish were mounted in the 3D-printed fish holder, secured with 3D-printed anchors, and the reservoir was filled with 1x Tricaine. Once the fish was secured, it was placed on a digital scale (SEAUMOON Digital Kitchen Scale) under the stereoscope. A micromanipulator was used to move the 3D-printed adaptor, align the ultrasonic scaler’s tip, and place 3g of pressure on the visible junction of the optic tecta and cerebellum directly over the confluence of sinuses. The ultrasonic scaler was activated for 1s with a frequency of 28kHz ± 3kHz at a power of 3 W (min dial setting), 6.4 W (3 dial setting), 9.8 W (5 dial setting), or 13.2 W (7 dial setting). Throughout the paper, the sham injury consisted of animals being anesthetized and placed in the fish holder with the dental tool placed on the skull for 3 seconds at 3 g of pressure, without any sonication. CVI and SHAM procedures were always conducted between 2 and 4 pm. All experiments have both sexes represented equally. After injury or sham injury, fish were revived with 80°C system water and observed for 3 hours before being transferred to the zebrafish system.

### Longitudinal, live confocal imaging

Adult zebrafish that were longitudinally images we individually housed in 0.8L tanks (Aquaneering ZT080) with a plant for enrichment. Fish were imaged before CVI, and 1-, 3-, 7-, 14-, and 21-dpi. On days fish were live imaged, food was withheld. Intubation of adult zebrafish and imaging of the meninges was conducted as described previously (Castranova et al., 2022; Castranova et al., 2021; Greenspan et al., 2024; Xu et al., 2015). In short, 126 mg/L Tricane in system water was placed in a 1L reservoir with an air stone, and a peristaltic pump (World Precision Instruments Peri-Star Pro PERIPRO −4 l) circulates the water first through a pulse regulator, then to the fish, and then back to the reservoir. The water is kept at 80°C by wrapping the silicon tubing (Tygon) around a heat block (SH100 Mini Dry Bath Hot Block). The imaging chambers were designed for either inverted or upright microscopes (details below) and the fish were held in place by sponges (anchored by magnets for the upright scope). Fish were imaged for up to 16 hours before being revived with 80°C system water and observed for 3 hours before transferred to the zebrafish system.

### CVI hemorrhage quantification

For quantification of hemorrhage area, fish lacking melanophores *nacre-/-,* were imaged with Leica (LEICA DFC7000 T) stereomicroscope 1 hour after injury to ensure hemorrhage into the meningeal space had finished. Using ImageJ, the skull area region of interest (ROI) was drawn around the parietal bone skull plates, whose edges are clearly visible and iridescent in the light images. Then the hemorrhage ROI was drawn around the red blood visible under the transparent skull. The percent area of the hemorrhage was calculated as (hemorrhage ROI area)/(skull plate ROI area)*100 and plotted as a proportion of skull area that had hemorrhage as either 0-25%, 25-50%, 50-75%, or 75-100%.

### CVI skull stain and quantification

For fluorescence imaging of the zebrafish skull, live animals were placed in system water with 0.01% Alizarin for 15 min as described previously (Bensimon-Brito et al., 2016). Fish were then rinsed 4 times in fresh system water for 30 minutes each for residual stain to leave the skin before being live imaged as described above. Alizarin Red S (Sigma Aldrich A5533) was kept as a stock solution of 0.5% pH 7.0 before dilution into system water. The skull has natural autofluorescence in the 405 nm and can be easily captured by boosting the laser power level according to the objective. On the Nikon Ti2 inverted microscope with Yokogawa CSU-W1 spinning disk confocal, Hamamatsu Orca Flash 4 v3 camera, with 20x water immersion 0.95 NA, the power for 405 nm = 95% and for 561 = 98%. Fish were live imaged with the intubation before and after injury and images were then denoised with NIS Elements AR 5.42.04 software using ‘NIS.ai’ and scored blindly. The scoring rubric (examples in Supp. Fig. 2F-Z) is as follows: Score 0 is an undamaged skull with all alizarin fluorescence intact. Score 1 has the alizarin stain scraped off the top of the skull. This is likely partially due to the short time of dosing live fish with the alizarin dye meaning that it did not penetrate the full thickness of the skull, leaving surface impacts very easy to visualize. Score 2 had some small surface level cracks inside of the scratched off alizarin stain that did not extend into the skull where the autofluorescence was intact. Score 3 the cracks became more substantial and in some cases extent outside of the alizarin absent divot. Cracks for score 3 were never full thickness through the skull which was easily visualized with the autofluorescence. Score 4 cracks visible with the alizarin correspond with a full thickness break in the autofluorescence through the skull. For all injuries was there was no visible leakage from inside the skull to outside by observation with a stereoscope.

### Fluorescence microscopy

Animals were euthanized according to the animal protocol via ice bath at 1-, 3-, 7-, 14-, and 21-days after CVI or 1 day after Sham injury (explained above). Fish were decapitated and placed in methanol free 4% paraformaldehyde (Electron Microscopy Solutions CAT #15710) buffered to 7.4 pH with 60mM HEPES overnight (not more than 16 hours) at 4°C on a rocker (about 12 full tilts per minute). After fixation, samples were rinsed twice for 5 minutes in PBS, followed by 5 days in 10% EDTA (pH 7.4) at room temperature. Samples were then rinsed for 24h in PBS without magnesium or chloride ions (GibocTM CAT #70011044) and sectioned into (150um-300um) sagittal slices with a Vibrating-feather blade on a microtome (Leica VT1000 S). Sections were stored in PBS at 4°C until staining.

For Edu staining, 24 hours before euthanasia 4ul of 10mM EdU was intraperitoneally injected as described previously (Lindsey et al., 2018). Vibratome sections underwent EdU staining following the manufacturer’s instructions (Click-iT EdU; Invitrogen, cat# C10340). In short, tissue sections were permeabilized for 20-minute incubation at room temperature in a permeabilization buffer (1% Bovine Serum Albumin (Thermo Fischer CAT #B14) and 0.5% Triton X-100 (Promega CAT #H5142) in PBS). Samples were washed twice for 5 minutes in 1x PBS with 1% BSA before incubation with the EdU solution at room temperature for 1 hour. Sections were washed twice for 5 minutes in 1X PBS solution with 1% BSA, counterstained with DAPI (1:10,000 (Thermo Fisher CAT #D1306)) and cleared with a clearing solution (1ml glycerol, 6ml 2,2’-Thiodiethanol, 3ml 1X PBS) (Costantini et al., 2015; Taimatsu et al., 2025) for 24h at 4°C on a slow rocker. Samples were mounted in clearing solution before imaging.

For TUNEL staining, tissue sections were permeabilized with 20μg/ml proteinase K in PBS for 5 minutes on a circular rotating table at 50 rpm before 2x 5-minute washes in PBS. Sections were post-fixed in 4% paraformaldehyde for 15 minutes and washed 5 times for 5 minutes with PBST (PBS with 0.01% Triton X-100). Afterwards, samples were further permeabilized with a 10-minute incubation in a permeabilization buffer (0.1% Triton X-100 and 1% sodium citrate (Thermo Fischer CAT #J63888.AK) in PBS). Sections were washed twice for 5 minutes with PBST and stained with TUNEL using kit instructions (TUNEL Assay Apoptosis Detection Kit, Biotium CAT #30074). Samples were incubated with TUNEL Reaction mix, which included 2.5 μL of TdT Enzyme to 50μL of TUNEL Reaction Buffer and DAPI for one hour at 37°C on a rocker in the dark. Sections were washed twice with PBST for 5 minutes before DAPI counterstain and cleared with LUCID.

Imaging of sections was done on a Nikon FNSP upright microscope with AXR scanning confocal. Images were acquired with a 20x water immersion objective (1.0 NA) with sequential resonance bidirectional mode with 8x averaging per channel, 1024×1024 pixel ratio, and 1.14 zoom using the Nikon Elements Advanced Research software. 405nm, 488nm, 561nm, and 640nm laser lines were used with an emission range of 429-474nm, 499-551nm, 571-625nm, and 662-737nm, respectively. Z-series stacks were captured with a <u>1.5-μm</u> step size and were processed, denoised, and stitched with Nikon Elements. A Maximum Intensity Projected image of 21 sections was imported to ImageJ for cell count image analysis. To quantify EdU^+^ or TUNEL^+^ cells, regions of interest were drawn around the dura, choroid plexus, leptomeninges, and superior cerebellum adjacent to injury using the Fiji polygon tool. Fiji Auto Threshold v1.18.0 Shanbag or Yen methods for Edu and TUNEL respectively. Cells were quantified using the Analyze Particles feature in Fiji with a set size of 3μm^2^-Infinity for EdU^+^ cells and 1μm^2^-Infinity for TUNEL^+^ cells.

### Behavior

Custom-designed infrared (IR) permeable-plastic tanks (Precision Plastics Inc.) were used for the diving tank assay as previously described (Anwer et al., 2021). These tall tanks (7 cm width, 52 cm height, 10.5 cm length) were designed with Infrared Transmitting Acrylic, making them impervious to visible light but not IR. Using the ZebraTower platform (Viewpoint Behavior Technology), 4 tanks faced the IR camera (maximum 30fps) with live tracking 2D sideview ZebraLab software (Viewpoint Behavioral Technology) to track and record locomotion. When running the assay CVI and SHAM treatments were every other animal to account for slight bench effects between runs as 2 animals were always CVI and 2 were SHAM for each run. 10-minute recordings were integrated into 5-minute periods. ‘Background refresh delay’ was set to 16 seconds, ‘detection sensitivity’ was set to 10, movement thresholds were set to 8 for ‘large’ and 4 for ‘inactivity’, and ‘X minimum size’ was set to 15. Each tank had 2 ROIs – one on the top half and one on the bottom. Adult Zebrafish were housed separate in 1.8L tanks at least 24h before diving tank assay. Before each experiment, fish were netted into a 600mL beaker with 200mL system water and kept in darkness for 10 minutes to avoid aversion behavior from netting and seeing researchers. Fish were then carefully poured into the behavior area for a total volume of 2 L of system water. After recording, each fish was transferred into their individual home tanks, and the IR tanks were cleaned with 70% ethanol and rinsed with system water between each run. Every fish was assayed once per time point, for a total of 6 assays per fish. All experiments were performed between 1:00 pm and 6:00 pm, or 5-10 hours into daylight cycle. As a positive control for anxiety response, a PLA-printed predatory toy (**Supp. File 13**) was used to chase the fish around their home tanks for 10 seconds before being put in the beaker.

For analysis of diving tank assay, the last 5 min integration period of the recording was used for % Time going slow, medium, or fast, the distance and velocity calculations, and % time in the top half of the diving tanks. Latency as the time it took the fish to enter the top half of the tank, was calculated by a researcher blindly recording the time after being transferred into the diving tank the fish settled at the floor of the tank and the time the animal crossed into the top ROI then calculating the seconds in between. As indicated above, slow speed was when the fish moved less than 4 cm/s, medium speed was between 4 and 8 cm/s, and fast was any speed greater than 8 cm/s. The tracks shown are also from the second 5-minute integration period as the calculations.

## 3D-cell live cell counts

Neutrophils were counted automatically using the Spots tool in IMARIS software. The cell diameter was set to 15μm for the XY plane and 30μm for the Z plane to account for Z-stretch during imaging and cell movement. Cells were tracked using the far-red channel with background subtraction enabled and counted within a restricted 3D region of interest (650μm Width, 650μm length, 110μm depth) around the area of injury. Neutrophil spots were manually removed if incorrectly assigned to debris or autofluorescence.

### Neutrophil tracking

Imaris Cell Tracking Software (Version 9.9.0) was used for semiautomated immune cell tracking. Confocal microscopy was used to generate fluorescent z-stack 10-minute movies which then were converted to 2-dimensional maximum intensity projections in Imaris. The “spots” tool was used for manual tracking of cells, with the “auto-connect to selected spot” setting enabled to facilitate continuous tracking. The total number of frames was divided by 10, and the movie was examined at each frame point, equally spaced, for a spot to follow from one side of the vessel to the other. Spots were manually tracked by every frame that they were seen. Computation of track speed and displacement was done by the Imaris distance-detection Matlab algorithms within the software. The color mapping of speed on Imaris tracks was done in the “color” tab, setting the color type to “statistics coded” for speed, and applying it to the tracks. Every image kept the colormap on the “spectrum” setting with a range of 0-150 for visualization of track speed.

### Quantification for vessel regrowth

To quantify vessel regrowth, a region of interest was made around the avascular area at every time point using the polygon tool in Fiji. The area of the region of interest was measured, and the percentage of vessel regrowth was calculated as described previously (Greenspan et al., 2024).

### Statistics

All statistics were done in Graphpad prism 10.0.2 (171) as described in the figure legends. * p < 0.05, ** p < 0.01, **** p < 0.0001.

### Data availability

All data are available upon request

### Arduino resources

Breadboard circuit- an image with legend showing how everything is connected Supplemental_16_Arduino circuit, code, components

## Acknowledgments

This research was supported [in part] by the Intramural Research Program of the National Institutes of Health (NIH). The contributions of the NIH author(s) are considered Works of the United States Government. The findings and conclusions presented in this paper are those of the author(s) and do not necessarily reflect the views of the NIH or the U.S. Department of Health and Human Services. The authors would like to thank the staff of the shared research facility for their dedication to the fish and ensuring researchers can depend on having healthy animals. They would also like to thank the Weinstein lab, Sheppard lab, NICHD Aquatics Affinity Group, and Ben Garcia who actively listened to presentations and offered critical discussion for the development of the presented project. This work was supported by the intramural program of the NICHD, NIH (ZIA-HD008915, ZIA-HD008808, ZIA-HD001011) to (BMW).

