## Supplemental Materials for "Live longitudinal imaging of meningeal cerebrovascular injury and its sequelae in adult zebrafish"

#### SUPPLEMENTAL FIGURE LEGENDS

**Supplemental Figure 1. Heterogeneity of the dural vasculature. A,** Pseudocolored confocal image of the dorsal head of a 6-month-old *Tg(kdrl:mcherry)*$^{y205}$ animal (same image as in Fig. 1C), annotated with the dural sinus vasculature pseudocolored in magenta and labeled, and the locations of other vessels in the image noted with other colors. **B-E,** Confocal images of the dorsal head of a 6-month-old *Tg(kdrl:mcherry)*$^{y205}$ shown at different magnifications, with the dural sinus vasculature pseudocolored in magenta in panel E. **F-S,** Confocal images of the dorsal heads of fourteen different 6-month-old *Tg(kdrl:mcherry)*$^{y205}$ siblings, revealing variation in the detailed pattern of the dural sinus vasculature (pseudocolored in magenta), despite similarity in their gross patterning. The bottom half of each panel shows a magnified portion of the image in the top half of the panel. Scale bars = 500 µm (A), 500 µm (B), 200 µm (C), 200 µm (D,E), 500 µm (top halves of F-S), 200 µm (bottom halves of F-S).

**Supplemental Figure 2. Circuit diagram for digital control of ultrasonic dental scaler.** This circuit replaces the foot pedal of an ultrasonic dental scaler, enabling precise, millisecond-level digital control of activation timing for use in a zebrafish injury model. The circuit consists of three components: an Arduino Uno microcontroller (A), a push-button module (B), and a 5V relay module (C). When the button (B) is pressed, it sends a 5V HIGH signal to the Arduino's (A) digital pin D2, allowing portable, standalone operation without a connected computer. The Arduino (A) then sends a 5V HIGH signal from digital pin D9 to activate the relay (C), which closes its normally open (NO) contacts and completes the circuit that powers the dental scaler. The relay (C) provides electrical isolation, enabling the low-voltage Arduino (A) to safely control the higher-voltage AC circuit used by the scaler.

**Supplemental Figure 3. Hemorrhage and effects on the skull after dural meningeal cerebrovascular injury. A-D,** Representative images of dural hemorrhage at 3 W (A), 6.4 W (B), 9.8 W (C), and 13.2 W (D) of power. Scale bars are 1.5mm. **E,** Schematic diagram of experimental workflow for Alizarin red staining, mCVI, and skull imaging. See Methods section for additional details. **F-Y,** Representative confocal images of alizarin red-stained skull surface bone (orange) and deeper autofluorescent skull bone (blue) on the dorsal skull of wild type adult zebrafish siblings before that were uninjured (F-I) or subjected to mCVI with 3 W (J-M), 6.4 W (N-Q), 9.8 W (R-U), or 13.2 W (V-Y) of power for 1 second. Images shown in panels are dorsal views 3D rendered to show "solid" views of the skull surface (F,J,N,R,V) or lateral views of a portion of the same confocal image stacks beginning from the lines on panels E and H, with alizarin + autofluorescence (G,K,O,S,W), alizarin alone (H,L,P,T,X), or autofluorescence alone (I,M,Q,U,Y). Scale bars = 50 µm. **Z,** Scoring rubric used to determine values graphed in Fig. 2M.

#### SUPPLEMENTAL MOVIE LEGENDS

**Supplemental Movie 1. Method of meningeal cerebrovascular injury.** A movie showing (1) an overview of the apparatus used to induce meningeal cerebrovascular injury (mCVI) in adult zebrafish, (2) higher magnification view of a hemorrhaging mCVI lesion being generated in the dural meninges of an adult fish, and (3) an adult fish swimming after mCVI injury.

**Supplemental Movie 2. Zebrafish skull remains intact after mCVI.** Overview schematic diagram and confocal images of alizarin-stained live adult zebrafish before and after mCVI at 6.4 W power. Alizarin labeling of the superficial skull (orange) and autofluorescence of the deeper skull (blue) shows that the skull remains intact after mCVI.

**Supplemental Movie 3. Novel tank diving test assay after mCVI.** Representative example video of a novel tank diving test to measure adult zebrafish anxiety. Tall test tanks (7 cm width, 52 cm height, 10.5 cm depth) opaque to visible light but transparent to infrared light were recorded with an infrared camera for 10 min (only 1-1/2 minutes shown). Fish occupancy of either the bottom or top of the tank was assessed in addition to other parameters. The time from when the fish settles on the bottom of the tank to when it first transitions to the top of the tank was recorded as the latency, an established measurement of anxiety. In the example shown here fish were tested at 3 days post-sham or mCVI injury,

**Supplemental Movie 4. Hematoma forms around dural venous sinuses after mCVI.** Transcranial confocal imaging of red blood cells stained with Hoechst nuclear dye (teal) flowing through the dural venous sinuses (magenta) of a *Tg(kdrl:mcherry)*$^{y205}$ transgenic adult zebrafish before (left) and after (right) mCVI. Hematoma is visible as stationary blood cells.

**Supplemental Movie 5. Vascular plasticity after mCVI.** Ten hour long live time-lapse transcranial confocal imaging of the dural venous sinuses (white) in uninjured (left) and 1 day post-mCVI (right) intubated adult *Tg(kdrl:mcherry)*$^{y205}$ transgenic adult zebrafish. Vessels are quiescent in the uninjured animal, but active angiogenesis is taking place in the vicinity of the injury in the animal with mCVI (example highlighted in teal).

**Supplemental Movie 6. Dural superior sagittal sinus blood flow is retained after mCVI.** Live transcranial confocal imaging of blood flow through the superior sagittal sinus (SSS) before (top) and immediately after (bottom) induction of mCVI at the nearby confluence of sinuses (COS) in an adult *Tg(kdrl:mcherry)*$^{y205}$ transgenic adult zebrafish (magenta) intravascularly injected with fluoromax beads (teal), showing that robust blood flow is maintainined through the SSS after COS mCVI.

**Supplemental Movie 7. Neutrophil dynamics in the injured dura immediately after mCVI.** Twelve hour long live time-lapse transcranial confocal imaging of neutrophils (magenta) in the dura of an intubated adult *Tg(lyz:DsRed2)*$^{nz50}$ transgenic animal, beginning immediately after mCVI injury. Neutrophils are initially sparsely distributed in the meninges but accumulate rapidly after injury, first rolling through the nearby dural sinuses and then accumulating in the meninges around the site of injury.

**Supplemental Movie 8. Neutrophils move slower through the dural sinuses after injury.** Ten-minute-long time lapse (only 6-9 minuts shown) of a single plane through the dura in a of an intubated adult *Tg(lyz:DsRed2)*$^{nz50}$ transgenic fish before and 1 day after mCVI. Commit tails show relative speed of neutrophis (Magenta) travelling through the superior sagittal sinus with red being fast and blue being slower. Neutrophils in the meningeal tissue were not tracked for vascular interactions for this study.

#### SUPPLEMENTAL FILES

**Supplemental File 1** CAD design for 3-D printed platform used to restrain adult fish for the mCVI procedure, designed with posts for securing bracing pins (Supp. File 12) and with a water reservoir for keeping the fish moist. Cavity width is 6mm at the top and 4mm at the base with a depth of 6mm.

**Supplemental File 2** CAD design for 3-D printed platform used to restrain adult fish for the mCVI procedure, designed with posts for securing bracing pins (Supp. File 12) and with a water reservoir for keeping the fish moist. Cavity width is 6mm at the top and 4mm at the base with a depth of 4mm.

**Supplemental File 3** CAD design for 3-D printed platform used to restrain adult fish for the mCVI procedure, designed with posts for securing bracing pins (Supp. File 12) and with a water reservoir for keeping the fish moist. Cavity width is 6mm at the top and 3mm at the base with a depth of 4mm.

**Supplemental File 4** CAD design for 3-D printed platform used to restrain adult fish for the mCVI procedure, designed with posts for securing bracing pins (Supp. File 12) and with a water reservoir for keeping the fish moist. Cavity width is 6mm at the top and 3mm at the base with a depth of 5mm.

**Supplemental File 5** CAD design for 3-D printed platform used to restrain adult fish for the mCVI procedure, designed with posts for securing bracing pins (Supp. File 12) and with a water reservoir for keeping the fish moist. Cavity width is 4.3mm at the top and 3.1mm at the base with a depth of 4.4mm.

**Supplemental File 6** CAD design for 3-D printed platform used to restrain adult fish for the mCVI procedure, designed with posts for securing bracing pins (Supp. File 12) and with a water reservoir for keeping the fish moist. Cavity width is 4.4mm at the top and 3.1mm at the base with a depth of 3.5mm.

**Supplemental File 7** CAD design for 3-D printed platform used to restrain adult fish for the mCVI procedure, designed with posts for securing bracing pins (Supp. File 12) and with a water reservoir for keeping the fish moist. Cavity width is 6mm at the top and 4mm at the base with a depth of 6.8mm.

**Supplemental File 8** CAD design for 3-D printed platform used to restrain adult fish for the mCVI procedure, designed with posts for securing bracing pins (Supp. File 12) and with a water reservoir for keeping the fish moist. Cavity width is 6mm at the top and 4.5mm at the base with a depth of 6.5mm.

**Supplemental File 9** CAD design for 3-D printed platform used to restrain adult fish for the mCVI procedure, designed with posts for securing bracing pins (Supp. File 12) and with a water reservoir for keeping the fish moist. Cavity width is 6mm at the top and 5mm at the base with a depth of 7mm.

**Supplemental File 10** CAD design for 3-D printed platform used to restrain adult fish for the mCVI procedure, designed with posts for securing bracing pins (Supp. File 12) and with a water reservoir for keeping the fish moist. Cavity width is 6mm at the top and 4.5mm at the base with a depth of 4.5mm.

**Supplemental File 11** CAD design for 3-D printed bracing pins used to secure adult fish onto the fish bed (Supp. Files 1-10) during injury and prevent movement while the fish is under anesthesia.

**Supplemental File 12** CAD design for 3-D printed adapter designed to hold the ultrasonic dental scaler during the mCVI procedure and mount the appliance onto a micromanipulator for precise control.

**Supplemental File 13** CAD design for 3-D printed novel object with predatory features at 20x the size of an adult zebrafish, used to chase fish for 30 seconds to induce anxiety as a positive control for the novel diving tank test.

**Supplemental File 14** Code used to program an Arduino UNO R4 Wifi for one-second activation of a Woodpecker UDS-K Ultrasonic Scaler.

### Supplemental Figure 1

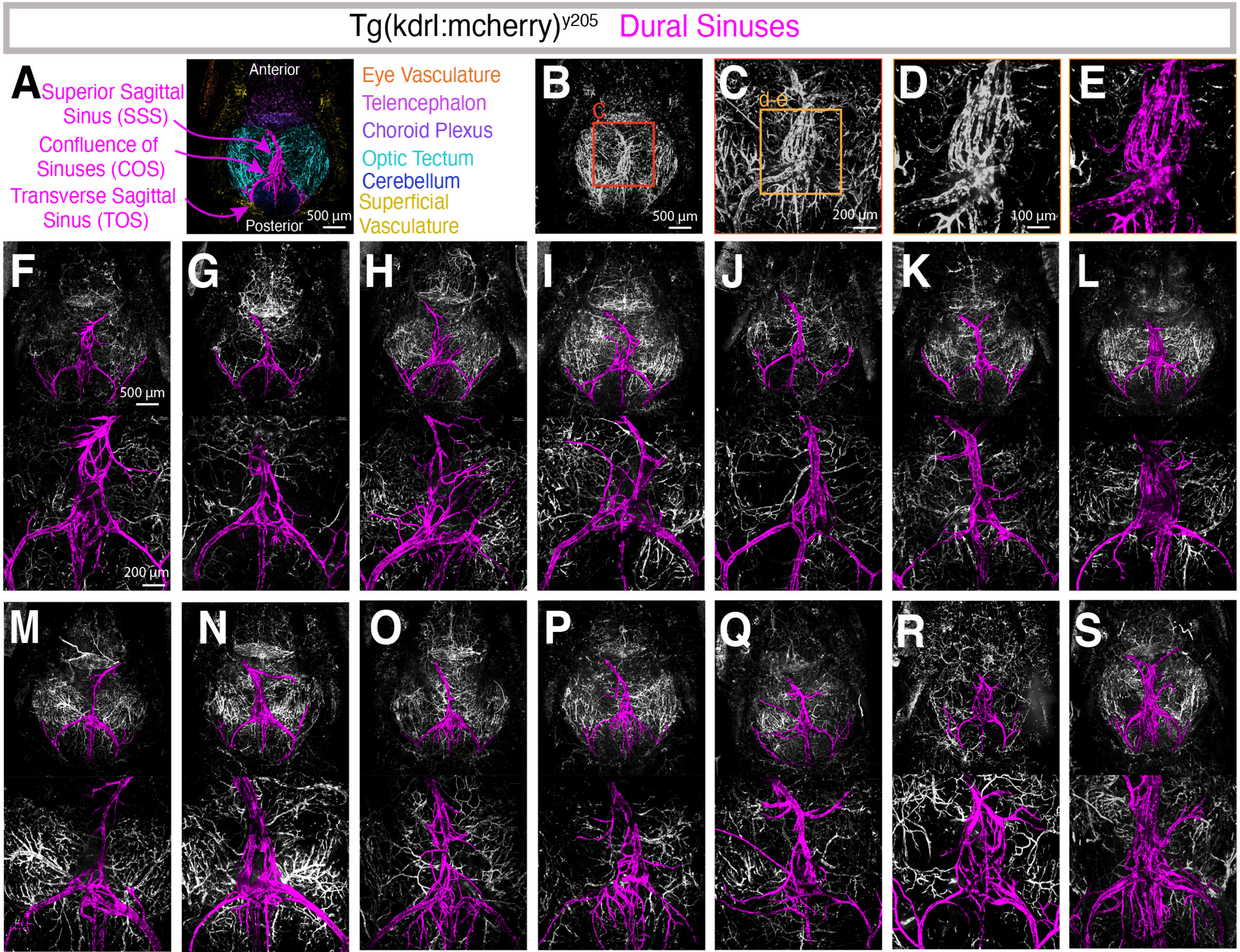

**Supplemental Figure 2**

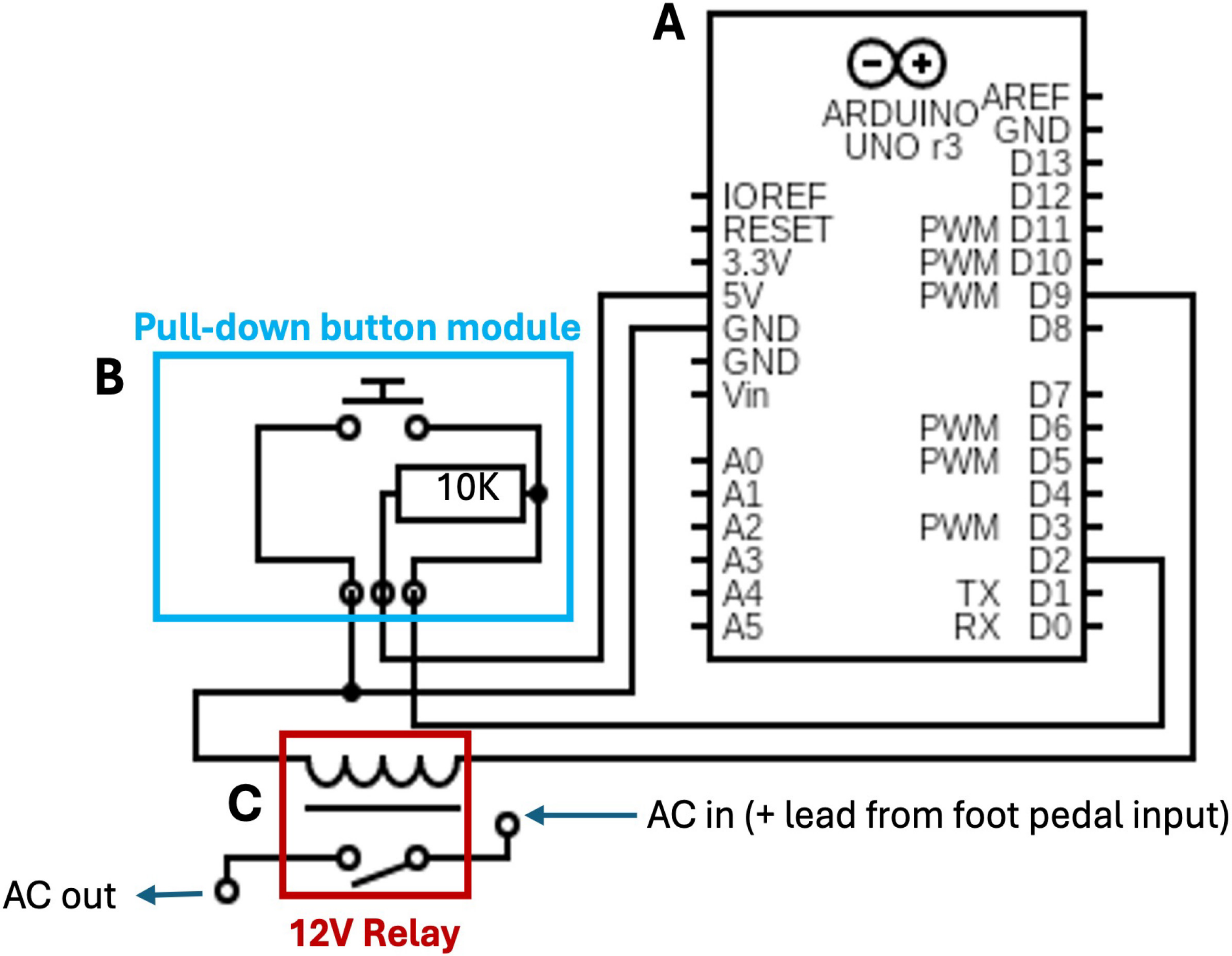

### Supplemental Figure 3

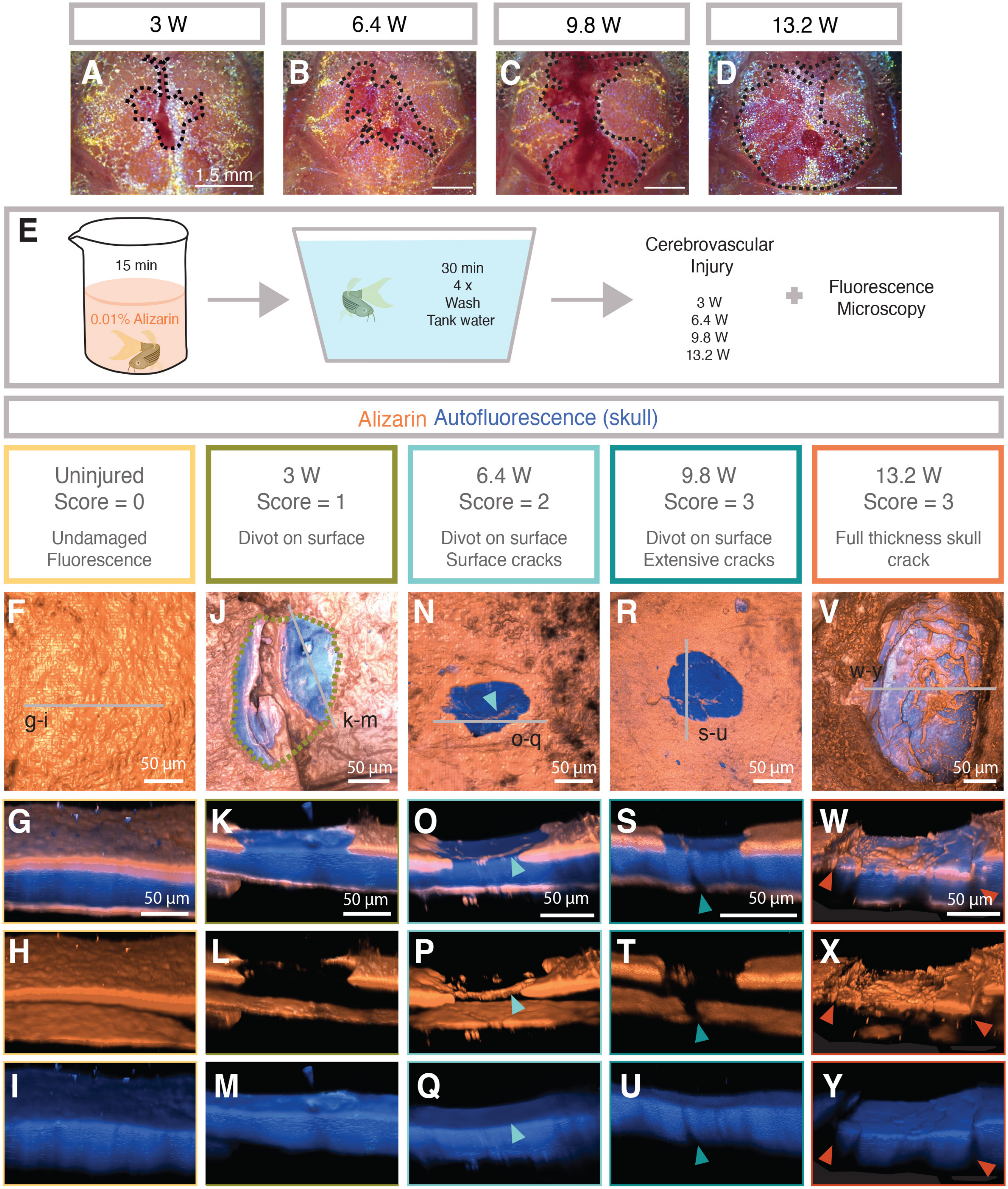


**Z**

| SCORE | DESCRIPTION |
|---|---|
| 0 | Undamaged Alizarin stain |
| 1 | Alizarin stain is scratched on the surface forming a divot |
| 2 | Alizarin divot + has superficial, small cracks |
| 3 | Large cracks are substantial and around alizarin divot |
| 4 | Fluorescence is discontinuous, Skull is broken |
