## Supplemental File 14 for "Live longitudinal imaging of meningeal cerebrovascular injury and its sequelae in adult zebrafish"

### Code Documentation

```
const int buttonPin = 2;        // Digital pin connected to the push
button
const int pedalPin = 9;         // Digital pin connected to the pedal

boolean pedalActivated = false; // Flag to track if the pedal has been
activated

void setup() {
  pinMode(buttonPin, INPUT_PULLUP);  // Set push button pin as input
with internal pull-up resistor
  pinMode(pedalPin, OUTPUT);         // Set pedal pin as output
}

void loop() {
  // Check if the push button is pressed
  if (digitalRead(buttonPin) == LOW && !pedalActivated) {
    // Turn on the pedal
    digitalWrite(pedalPin, HIGH);
    pedalActivated = true; // Set the flag to indicate that the pedal
has been activated
    delay(1000); // Wait for half a second (1000 milliseconds)
    // Turn off the pedal
    digitalWrite(pedalPin, LOW);
    pedalActivated = false;
  }
}
```

#### Code Explanation

- **buttonPin**: Configured as an input with an internal pull-up resistor. It reads the state of the push button.
- **pedalPin**: Configured as an output to control the relay.
- **pedalActivated**: A boolean flag to track the activation state of the pedal.
- **setup()**: Initializes the pin modes for the button and pedal.
- **loop()**: Continuously checks if the button is pressed. If pressed and the pedal is not already activated, it turns on the pedal for 1 second and then turns it off.
